# Characterization of the cardiac proteome of wild-type transthyretin amyloidosis cardiomyopathy

**DOI:** 10.1101/2025.09.29.679210

**Authors:** Marcus Rhodehamel, Vivek P. Jani, Ryan Gross, Krish Dewan, Atharva Mulay, Navid Koleini, Matt Foster, Kavita Sharma, M. Imran Aslam, Joban Vaishnav, Dawn E. Bowles, David A. Kass, Mark J. Ranek

**Author notes:** **Corresponding Author:** Mark J. Ranek, PhD, Department of Medicine, Division of Cardiology, The Johns Hopkins Medical Institutions, 720 Rutland Avenue, Ross 850, Baltimore, MD 21205.

## Abstract

**Introduction:** Myocardial accumulation of the protein transthyretin (TTR) can result in amyloid TTR cardiomyopathy (ATTR-CM), a form of restrictive heart disease with limited therapies and still generally poor clinical outcomes. The mechanisms by which TTR fibril accumulation elicits cardiac toxicity at the protein level remain largely unknown. Accordingly, we performed untargeted proteomics of ventricular myocardium from patients with ATTR-CM versus controls.

**Methods:** Myocardial tissue from non-failing (NF) controls (n=7) and ATTR-CM (n=4) were assayed by mass spectrometry. HFrEF, HCM, and HFpEF proteomics were acquired from published databases.

**Results:** A total of 539/7093 (7.6% of total) proteins were found to be differentially expressed in ATTR-CM, 227/359 (42%) upregulated and 312/539 (58%) downregulated. Gene ontology pathway analysis found that downregulated proteins were enriched for oxidative phosphorylation and mitochondrial protein translation pathways, while upregulated proteins were enriched for enhanced endocytosis and intracellular vesicle mediated transport. The latter is not observed in other forms of heart failure. We further identify a profound downregulation of sarcomere protein content, which is also not seen in other cardiomyopathies.

**Conclusion:** The ATTR-CM myocardial proteome identifies endocytosis and intracellular transport as uniquely upregulated processes, whereas sarcomere protein content is uniquely downregulated. Both maybe potential therapeutic targets.

## INTRODUCTION

Transthyretin (TTR) amyloid cardiomyopathy (ATTR-CM) is a form of restrictive cardiomyopathy characterized by infiltration and accumulation of TTR fibrils in the myocardium, that result in increased ventricular stiffness, hypertrophy, electrical conduction abnormalities, and marked diastolic dysfunction^1-3^. Clinical outcomes for patients with ATTR-CM are often poor, in part due to limited treatment options^4^. It is currently thought that the underlying disease pathogenesis involves dissociation of the tetrameric TTR complex into monomers and eventual amyloid fibril generation^2, 3^. Deposition of these TTR fibers increases diastolic stiffness and insulates and disrupts normal physiological conduction^5^. Disease modifying therapy involves stabilizing the monomeric form of the TTR protein preventing the formation of new fibrils. Early diagnosis and treatment with these therapies improves outcomes and quality of life in ATTR-CM patients, but they only slow progression of disease and do not reverse its course, in part because they cannot impact amyloid fibrils already present in the myocardium^2, 3, 6-8^. While recent studies have found amyloid fibrils are cytotoxic and can directly alter cardiomyocyte function^9-11^, little remains known about this process at molecular levels.

The majority of proteomic studies of ATTR-CM patients have been solely from blood, and data on the myocardial proteome remains limited^12-15^. Accordingly, this study aimed to profile the proteome of non-hereditary ATTR-CM (ATTRwt-CM) myocardium in comparison to non-failing controls. We find increased expression of proteins linked to endocytosis and vesicle-mediated transport, whereas expression of thick filament, mitochondrial protein translation, and ATP synthesis electron transport (OxPhos) proteins are reduced. The latter is also found in hypertrophic cardiomyopathy and heart failure with reduced [HFrEF] or preserved [HFpEF] ejection fraction^16-19^. Several of the endocytic and inflammatory processes identified, however, are unique to ATTR-CM, that could provide new insight into its underlying pathobiology.

## METHODS

### Data Availability

Analyzed proteomics files containing the individual protein abundance values along with the de-identified metadata are available in the online supplementary information. Python and R-scripts are available upon reasonable request.

### Study Population

Myocardial tissue was obtained from cardiectomized patients with ATTR-CM (n=4) and non-failing (NF) controls (n=7) at time of heart transplantation, left ventricular assist device implantation, or septal myectomy at the Duke University Hospital. NF controls were procured from deceased brain-dead organ donors whose hearts were not used for transplantation. Patient (or surrogate) consent was obtained for all samples included in the study. The study was approved by the Duke Human Heart Repository (DHHR) institutional review board (Pro00005621). Demographic and basic echocardiographic data were obtained prior to cardiectomy. Additional details are described elsewhere^20^.

### Myocardial Tissue Procurement and Processing

Explanted hearts were excised from patients or donors by surgical staff and fully submerged in cold PBS (VWR cat. 76371-734; 0°C) immediately upon removal. This protocol is described in detail elsewhere^20^. Briefly, the submerged hearts were then transported on ice to the dissection lab for further processing. On a chilled cutting board (4°C), a “bread-loaf” slice from the mid-ventricular region (encompassing both ventricles and septum) was isolated and dissected into left ventricular, right ventricular, and septal strips using a sterile surgical scalpel. Strips tissue from the left ventricle, right ventricle, and septal wall were further sectioned into 200 – 400mg pieces and flash-frozen in liquid nitrogen. Cryogenically preserved tissue samples were transferred to - 80°C freezers for long term storage prior to downstream assays. Excess cardiac tissue post sectioning was fixed in 10% neutral-buffered formalin and submitted for further histopathological medical examination.

### MS Data Processing and Curation

Raw MS tracings were quantified for protein abundance using Spectronaut’s integrated MaxLFQ algorithm^21^. Proteins that had ≥50% missing values across any group (non-failing, failing) were removed. Summary of excluded proteins due to insufficient detection is reported in **Supplemental Figure S3**. Proteins associated with blood contamination were removed as defined by manual curation and explicitly stated in the data supplement. Data for proteins that had ≤50% missing values were imputed using a standard MissForest (ver.2.5.5; missingpy ver. 0.2.0) in Python (ver. 3.10.16)^22^. Subsequent and analysis is based on imputed data values. MS proteomic repositories for HCM, HFrEF, and HFpEF were previously curated and sourced for publicly available archives^16, 17^. Raw and imputed protein abundance values are provided **Supplementary Figure S3**.

### Western Blot Analysis

Lysates from explanted ATTR-CM and non-failing control tissue were extracted by high-speed bead homogenization in lysis buffer (6M urea, 5% SDS, 1 mM DTT, 1X protease inhibitor [Sigma Aldrich], and 1X phosphatase inhibitor [phosSTOP, Roche]) and clarified by high-speed centrifugation (16,000 g for 1 minute). Protein concentration was assayed by a Bradford assay. Extracts were diluted with 1X Laemmli Buffer and loaded into Bio-Rad Criterion TGX Gels (4-20% gradient). Gels were transferred to nitrocellulose membranes. The following antibodies were used: Myosin Light Chain 2 (MYL2, Novus Biologicals, EPR3741), phosphoserine 15 MYL2 (Invitrogen PA5-104265), and phosphoserine 19 MYL2 (Rockland 600-401-416). Gels were imaged (Odyssey, Li-Cor) and band intensity quantified ImageStudio and normalized to total protein loaded (BioRad Total Protein Stain 520 nm).

### Statistical Analysis

Demographic and clinical data are provided as mean±SD for normally distributed variables or median (IQR) for those non-normally distributed. Log_2_ fold changes of protein expression vs. NF controls were calculated from imputed values, and differential expression was assessed using a two-tailed t-test with Benjamini-Hochberg correction to adjust for multiple hypothesis testing (adjusted p□<□0.05 was considered statistically significant). These were compared to previously published proteomic datasets for hypertrophic cardiomyopathy (HCM)^17^, HF with reduced ejection fraction (HFrEF)^17^, and HF with preserved ejection fraction (HFpEF)^16^. Proteins were cross-referenced and manually curated against the UniProt human protein database, and all proteins converted to standardized gene IDs. Principal component analysis (PCA) was performed on imputed values using the scikit-learn (ver. 1.6.1) package in Python. Principal component analysis (PCA) was performed on imputed values using the scikit-learn (ver. 1.6.1) package in Python. Detailed loadings for each of the proteins are provided in **Supplemental Figure S3**. Volcano plots were constructed plotting log_2_ fold changes against -log_10_ adjusted p-values. Gene Ontology (GO) pathway analysis was performed in R (ver. 4.3.2) using the enrichGO function from clusterProfiler library (ver. 4.10.1) and the org.Hs.eg.db (ver. 3.18.0) for the database reference. Gene Ontology (GO) pathway analysis was performed in R (ver. 4.3.2) using the enrichGO function from clusterProfiler library (ver. 4.10.1) and the org.Hs.eg.db (ver. 3.18.0) for the database reference for differentially expressed proteins. GO pathways were curated using a Jaccard index to minimize pathway redundancy (**Supplementary Figure S3**). For all analyses, the threshold for significance was an adjusted P < 0.05, or unadjusted P < 0.05 for previously published datasets.

## RESULTS

### Patient Characteristics

Clinical characteristics of ATTR patients are included in **Table 1**. Age and sex were matched for the two patient groups. Invasive hemodynamics were consistent with end-stage ATTR-CM: right atrial pressure (21 ± 5 mmHg), mean pulmonary artery pressure (39 ± 6 mmHg), and pulmonary capillary wedge pressure (29 ± 5 mmHg), pulmonary vascular resistance (3.1 ± 1.7 Wood Units); all markedly increased versus normal. Cardiac index (1.6 ± 0.2 L/min/m^2^) was markedly reduced. On echocardiography n=1 of 4 ATTR-CM patients had EF < 40%. The remaining relevant clinical hemodynamic parameters are summarized in **Table 2**.

**Table 1.**
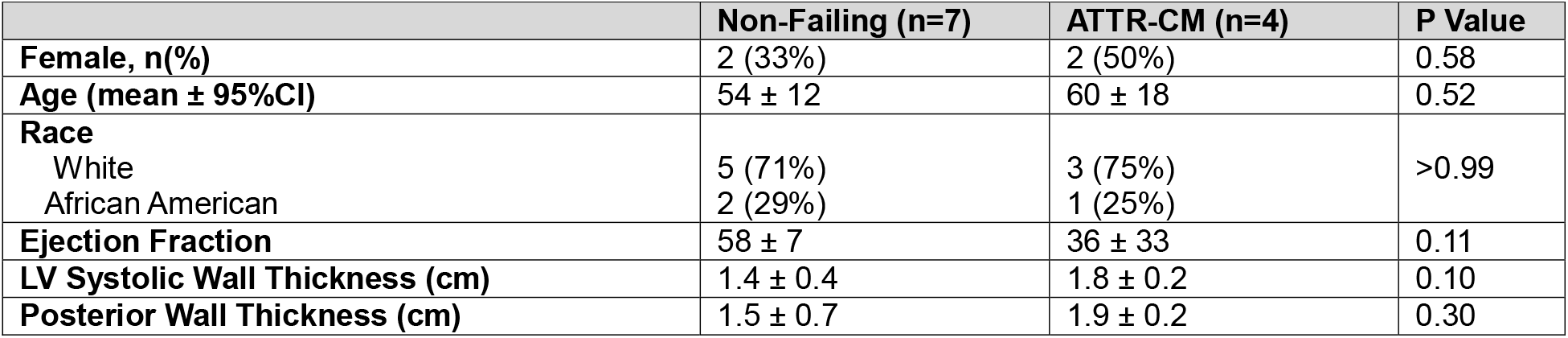
Clinical Characteristics of Non-failing Control and ATTR-CM Patients Included in MS Proteomics Analysis. P values are from a Mann-Whitney U test. LV – Left Ventricular

**Table 2.** Clinical hemodynamics of ATTR-CM patients.

|  |  |
| --- | --- |
| Heart Rate (bpm) | 87 ± 14 |
| Right Atrial Pressure (mmHg) | 21 ± 5 |
| Right Ventricular Systolic Pressure (mmHg) | 52 ± 6 |
| Right Ventricular Diastolic Pressure (mmHg) | 16 ± 7 |
| Mean Pulmonary Artery Pressure (mmHg) | 39 ± 6 |
| Pulmonary Capillary Wedge Pressure (mmHg) | 29 ± 5 |
| Pulmonary Vascular Resistance (Wood units) | 3.1 ± 1.7 |
| Cardiac Output (L/min) | 3.3 ± 0.7 |
| Cardiac Index (L/min/m<sup>2</sup>) | 1.6 ± 0.2 |
| Arteriovenous Oxygen Gradient (mL/dL) | 7.7 ± 0.7 |

### Differential Expression Analysis of the ATTR-CM Proteome

A total of 7093 unique proteins were identified in the proteome. Principal component analysis of these data is shown in **Figure 1A**. Notably, NF and ATTR-CM proteomes showed marked separation along the first principal component. Individual loadings for the top 20 relatively increased or reduced proteins defining a greater PC_1_ vector are identified in **Figure 1C**. Those higher in ATTR-CM were associated with nucleoside and phospholipid catabolism and transport, while those reduced were related to mitochondrial electron transport and oxidative phosphorylation.

**Figure 1.**
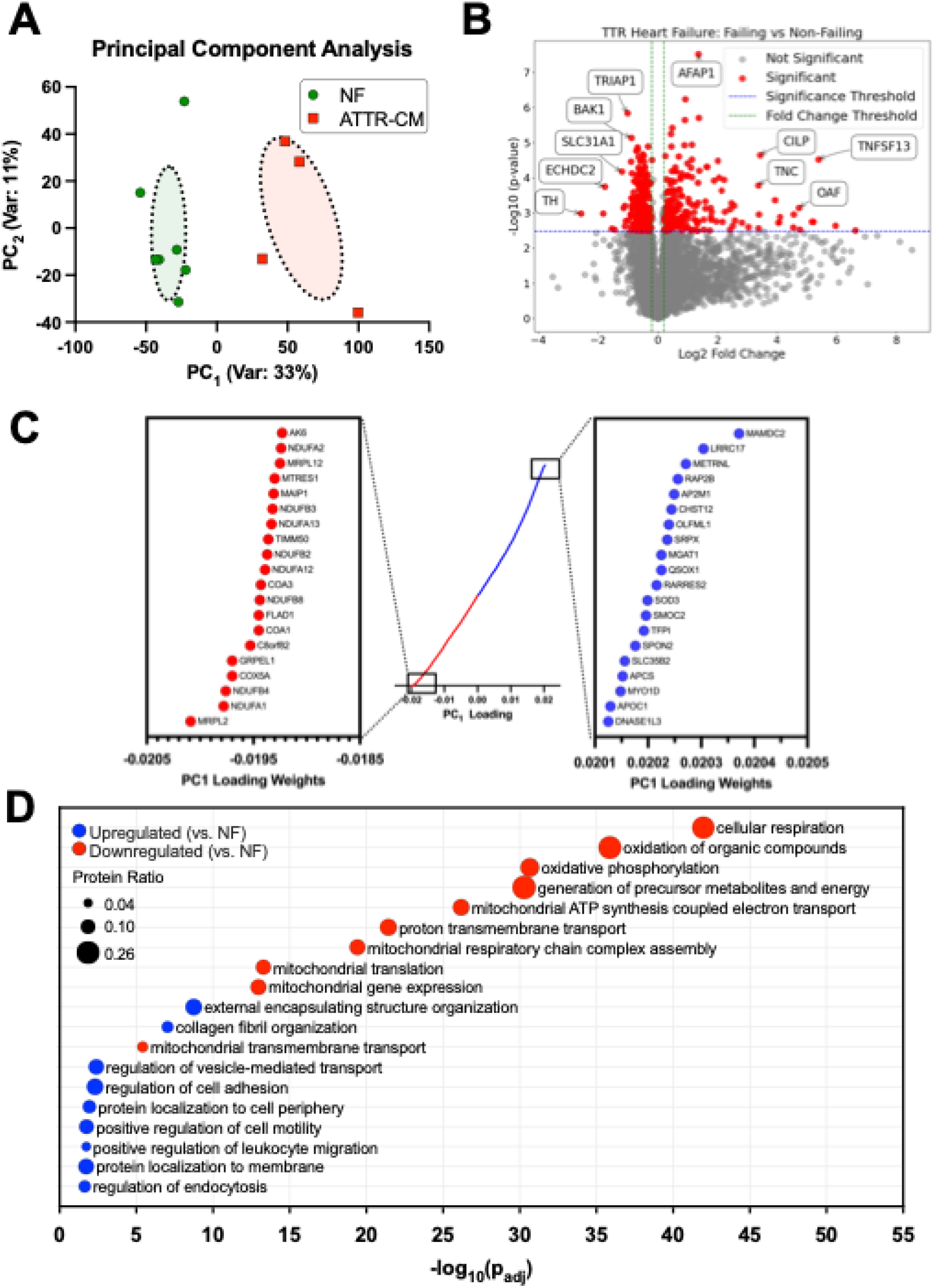
**(A)** Two-dimensional principal component analysis (PCA) of MS-based proteomics from 4 failing ATTRwt-CM myocardium (red) and 7 non-failing controls (green). **(B)** Volcano plot (log_2_ fold change vs. –log_10_ *p*-value) showing differentially expressed proteins in ATTRwt-CM versus controls. The top five upregulated and downregulated proteins are labeled based on magnitude of differential expression and statistical significance. **(C)** Dot plot of the full protein spectrum for PCA loading weights on PC1 (center), with expanded views of the top 20 most impactful positive (right) and negative (left) contributors to PC1. **(D)** Gene Ontology: Biological Process (GO:BP) enrichment analysis of significantly upregulated (red) and downregulated (blue) pathways (*Padj* < 0.05). The x-axis is the log transformation of the adjusted p-value. Circle size reflects protein ratio, which corresponds to the proportion of identified differentially expressed proteins to all known proteins in the pathway. Representative pathways were selected using Jaccard indexing to reduce redundancy and enhance interpretability, full list of proteins identified in **S3**. Color coding represents *P* value after Benjamini-Hochberg for multiple hypothesis comparison.

Differential expression analysis of the ATTR-CM vs. NF proteome (summary volcano plot shown in **Figure 1B**) identified 539/7093 (7.6% of total) significantly differentially expression proteins. Of these 227 (42%) were upregulated in ATTR-CM, while 312 (58%) were downregulated vs. NF controls. Gene ontology enrichment analysis (**Figure 1D**) found upregulated proteins enriched for endocytosis, intracellular vesicle mediated transport, cell motility, protein localization, and immune cell migration processes, whereas downregulated proteins were enriched for oxidative phosphorylation, mitochondrial protein translation, mitochondrial gene expression, and impaired transmembrane transport. Key proteins associated with metabolic dysfunction are shown in **Table 3**. A full list of all differentially expression proteins is provided in **Supplemental Figure S3**.

### Reduced Sarcomere Protein Expression in ATTR-CM

ATTR-CM is largely considered to be a disease of extracellular deposition of TTR amyloid fibrils. However, the disease also presents with intracellular cardiomyocyte abnormalities, with one recent study finding reduced sarcomere contractility and hypo-phosphorylation of cardiac troponin I and myosin binding protein C^9, 23^. Accordingly, we queried the proteome to better identify sarcomeric defects. Targeted analysis of sarcomere proteins is shown in **Figure 2A**. Surprisingly, the expression of beta myosin heavy chain (p=0.006), cardiac myosin binding protein C (p=0.006), tropomyosin (p=0.02), and all subunits in the troponin complex (p=0.006 for all) were reduced in ATTR-CM vs. NF controls. Western blot validation of these findings by total protein stain from ATTR-CM tissue lysates is shown in **Figure 2B** (raw gels in **Supplemental Figure S1-2**,**7**). In heart failure and during pathologic hypertrophy, the myosin heavy chain isoform shifts from Ill (faster) to β (slower), which is quantified by an increase in β/Ill myosin isoform ratio^24^. However, unlike other cardio-myopathies, we found no difference in β/Ill myosin isoform ratio in ATTR-CM vs. NF controls. These findings are unlikely to be explained by an increase a proportionate increase in TTR as cardiac phospholamban (p=0.65), also known to be enriched in cardiomyocytes vs. other cell types), was not different between ATTR-CM and NF controls. This suggests that the downregulation of sarcomeric proteins is specific, rather than a generalized reduction caused by the increased presence of extracellular matrix proteins.

**Figure 2.**
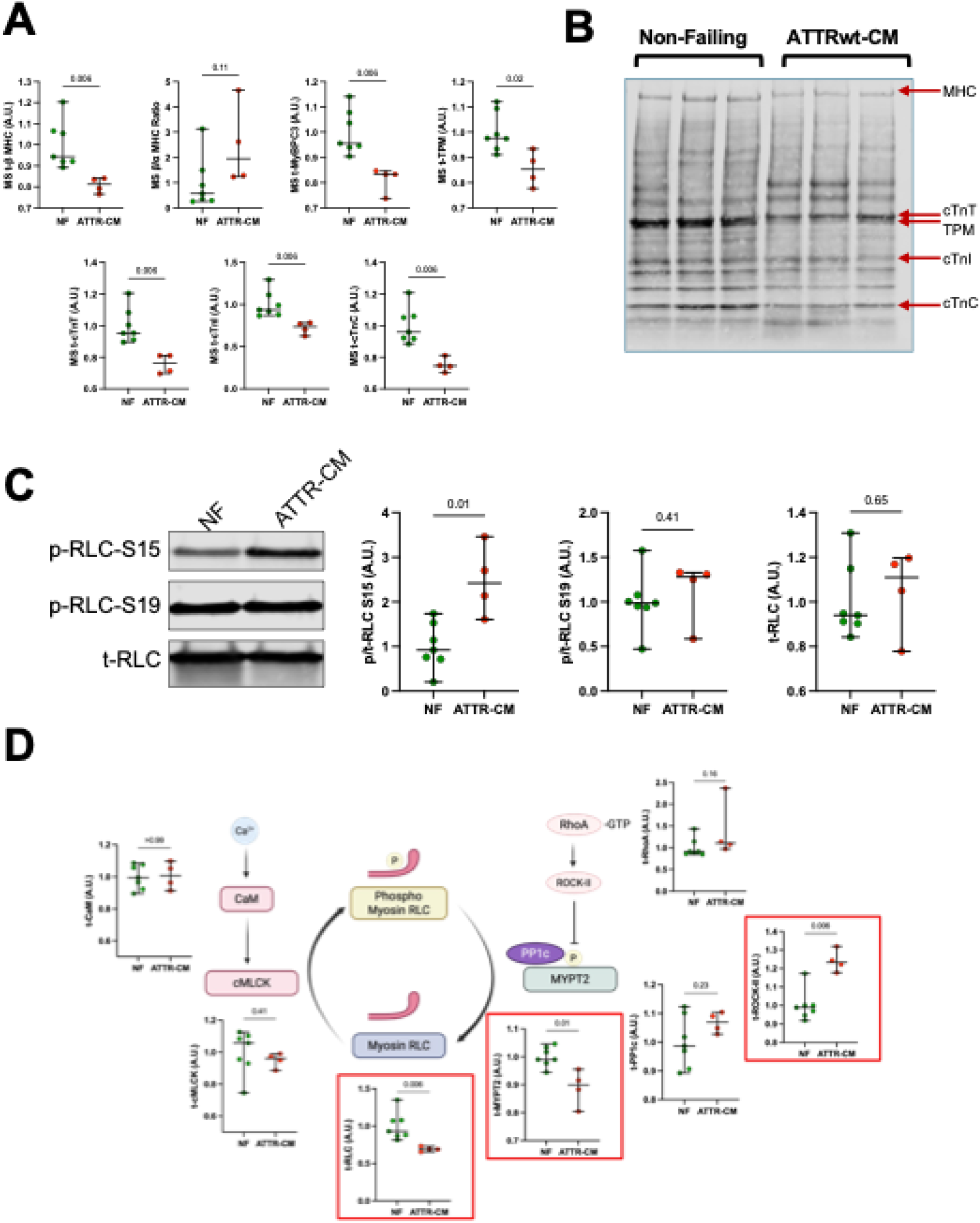
Sarcomere Proteins Expression is Reduced and RLC S15 Phosphorylation Increased in ATTRwt-CM: **(A)** Mass spectrometry abundance of contractile proteins: beta myosin heavy chain (β MHC), ratio of beta to alpha myosin isoform (β/α MHC), myosin binding protein C (MYBPC), tropomyosin (TPM), cardiac troponin T (cTnT), cardiac troponin I (cTnI), and cardiac troponin C (cTNC) showing reduced expression in ATTRwt-CM compared to non-failing myocardium. **(B)** Total protein stain western blot of myocardium lysates from ATTRwt-CM and non-failing controls, probed for sarcomere proteins, validating reduced contractile protein expression. **(C)** Representative western blot and quantification of S15 and S19 regulatory light chain (RLC) phosphorylation in ATTRwt-CM, demonstrating increased S15 phosphorylation. (**D)** Mass spectrometry–based abundance of proteins in the ROCKII–RLC signaling pathway in ATTRwt-CM: calmodulin (CaM), cardiac myosin light chain kinase (cMLCK), regulatory light chain (RLC), regulatory subunit of myosin light chain phosphatase (MYPT2), ras homolog family member A (RhoA), protein phosphatase 1 catalytic subunit (PP1c), and rho-associated coiled-coil-containing protein kinase 2 (ROCKII) showing increased ROCKII expression with reduced total RLC and MYPT2.

Alterations in sarcomere protein expression per se do necessarily imply functional depression, as their function often depends on post-translational modifications^25^. Using our differential expression analysis, we sought to identify other plausible mechanisms of sarcomere dysfunction related to sarcomere protein post-translational modification, specifically phosphorylation. Among upregulated pathways, seven were associated with serine/threonine kinase function and regulation, among them Rho-associated coiled-coil containing protein kinase 2 (ROCK-II). In the sarcomere, increased ROCK-II is often associated with phosphorylation of the regulatory light chain (RLC, gene MYL2)^26, 27^. Immunoblot of the two key phosphorylation sites on RLC, serine 15 (S15) and 19 (S19), revealed a 2.4-fold increase in phosphorylation of S15 (p=0.01) in ATTR-CM vs. NF (raw gels; **Supplementary Figure S1-2**). No change was observed in RLC S19 phosphorylation (p=0.41). The ATTR-CM proteome associated with the ROCK-II-mediated phosphorylation of RLC was investigated to identify perturbations in protein regulation. This analysis showed a reduction in abundance of both RLC (p=0.006) and primary phosphatase-scaffolding subunits responsible for RLC S15 dephosphorylation, myosin phosphatase 2 (MYPT2, p=0.01), in ATTR-CM vs. NF controls (**Figure 2C**).

### Comparisons of ATTR-CM to other Cardiomyopathies

To identify pathways uniquely dysregulated in ATTR-CM compared to other inherited and acquired cardiomyopathies (HCM, HFrEF, and HFpEF), data from previously reported cardiac proteomes from these syndromes were examined^16, 17^ (**Figure 3**). Pathways associated with protein import, endocytosis, vesicle mediated transport, and cellular motility were upregulated in ATTR-CM compared to all three other syndromes (**Supplementary Figures S4-6**). HCM, HFrEF, and HFpEF, all had reduction in proteins involving mitochondrial translation, mitochondrial gene expression, and NADH reduction pathways of the electron (**Supplementary Figures 4-6**). In the ATTR-CM proteome, there was even greater downregulation of these pathways than that in HFrEF, HFpEF, and HCM.

**Figure 3.**
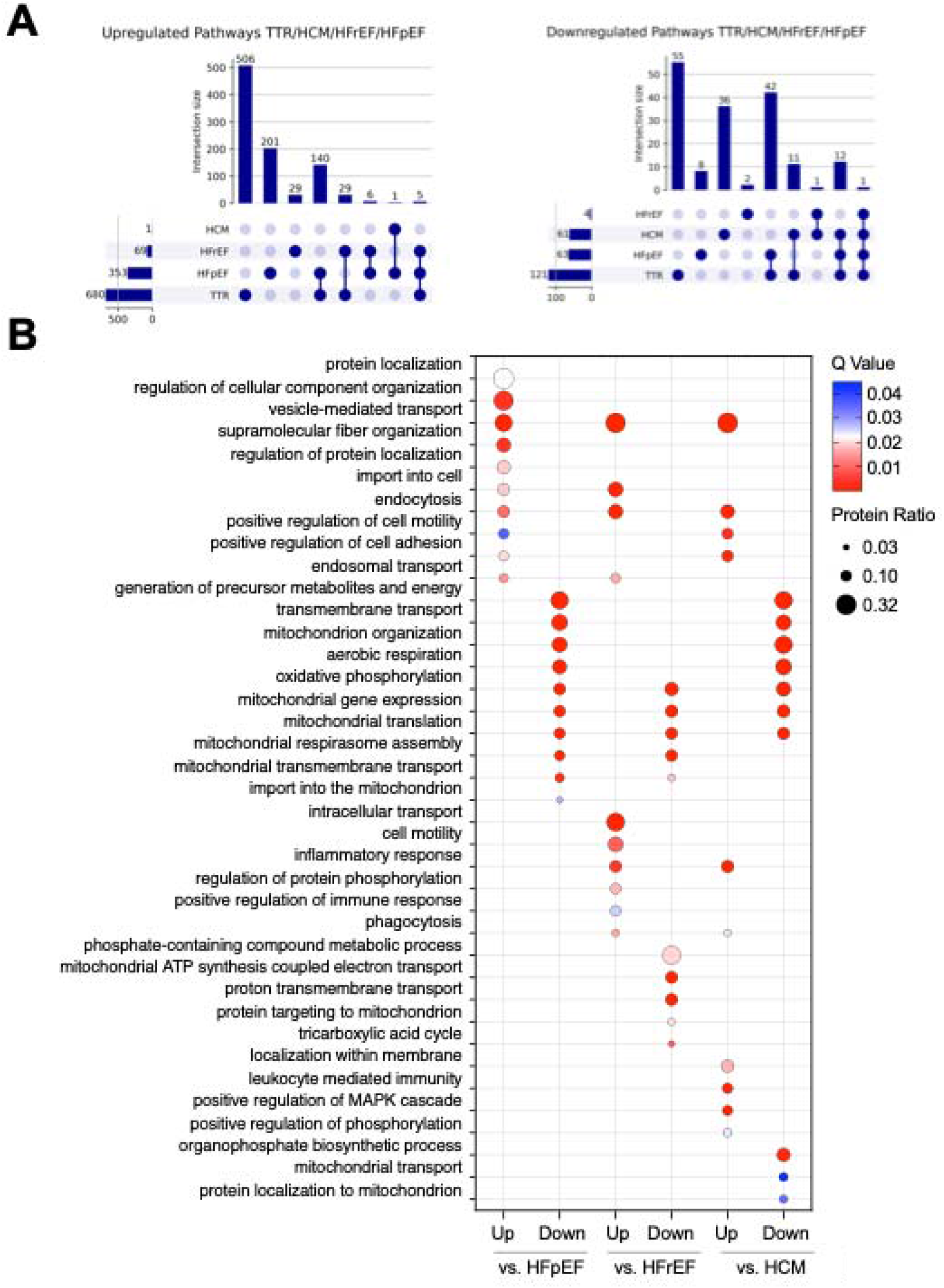
**(A)** UpSet intersection plot of cross-disease pathway distribution in ATTRwt-CM, HCM, HFrEF, and HFpEF, showing greatest overlap between ATTRwt-CM and HFpEF. Vertical bars indicate the number of pathways for each intersection; selected pathways for each combination are listed below the plot. **(B)** Gene Ontology: Biological Process (GO:BP) enrichment analysis comparing significantly upregulated and downregulated pathways (*P-value* < 0.05) in ATTRwt-cm vs HCM, HFrEF, and HFpEF. The x-axis is the log transformation of the adjusted p-value. Circle size reflects protein ratio, which corresponds to the proportion of identified differentially expressed proteins to all known proteins in the pathway. Representative pathways were selected using Jaccard indexing to reduce redundancy and enhance interpretability, full list of proteins identified in **S4-6**. Color coding represents *P* value.

Upon comparison of pathways identified as significantly upregulated in ATTR-CM, 506/680 (74%) were uniquely upregulated in ATTR-CM but not in HCM, HFpEF, or HFrEF. 145/353 (41%) pathways and 34/69 (49%) were similarly upregulated in ATTR-CM vs. HFpEF and HFrEF, respectively. Only one shared upregulated pathway was observed in ATTR-CM and HCM. 55/121 (45%) were uniquely downregulated in ATTR-CM and not observed in HFpEF, HFrEF, or HCM. HFpEF and ATTR-CM shared 42/63 (67%) downregulated pathways, while HCM and ATTR-CM shared 24/61 (39%) downregulated pathways. No downregulated pathways in ATTR-CM were also downregulated in HFrEF.

## DISCUSSION

The present proteomics analysis of explanted ventricular myocardium from ATTR-CM patients vs non-failing controls reveals several important findings. First, endocytosis and vesicle trafficking pathways are upregulated in ATTR-CM vs. NF controls. These pathways are unique to ATTR-CM, e.g. not found in the other cardiomyopathies investigated (i.e., HFpEF, HFrEF, HCM)^12, 17, 28^. Moreover, like many other forms of heart failure and inherited cardiomyopathies, ATTR-CM myocardium exhibits downregulation of mitochondrial and oxidative phosphorylation proteins as well as proteins related to mitochondrial protein transcription and translation. Notably, the degree of downregulation is greater in ATTR-CM. We also find significantly less sarcomere protein expression levels. This could reflect dilution of these normally highly abundant proteins due to the deposition of a substantial amount of TTR. These results highlight dysfunction of several key pathways, mostly related to endocytosis, vesicular trafficking and sarcomere protein expression, that are unique to ATTR-CM and may serve as novel therapeutic targets.

The majority of proteomic studies in ATTR-CM have been obtained from serum or mass spectrometry-based phenotyping of the protein composition of amyloid plaques themselves^13-15^. Serum from ATTR-CM patients has been observed to be enriched for proteins associated with extracellular transport and inflammation^15^. These results are somewhat nonspecific and provide little insight into myocardial ATTR-CM pathophysiology. Deep profiling of cardiac amyloid plaques have identified upregulation of signature amyloidosis protein (e.g., APOE, APOA4, and SAP) as well have unique proteomic profiles associated with ATTR and immunoglobulin light chain amyloidosis (AL), the other major etiology of cardiac amyloidosis^13,14^. The latter was a study performed in 292 patients ATTR patients and 139 AL patients and found upregulation of complement proteins in ATTR and keratin in AL^13, 14^. Notably, this study found autophagy to be a unique pathway downregulated in ATTR amyloid plaques^13^. Our study identified a significant upregulation of endocytotic pathways, which, when considered with reduced autophagy, may provide a plausible mechanism for the pathogenicity of extracellular amyloid fibrils.

We are only aware of one other study that performed proteomic profiling of myocardial tissue from 76 ATTR-CM patient endomyocardial biopsies compared to the proteome from 27 AL amyloidosis patient biopsies^14^. ATTR-CM was associated with downregulated metabolic pathways, consistent with our study, whereas AL amyloid samples were enriched for endocytotic and ribosomal proteins. This study did not have a non-failing control group, but did analyze biopsies rather than explanted hearts, so potentially assayed less terminal disease.

One intriguing finding in our study was upregulation of endocytotic and vesicular trafficking proteins in ATTR-CM vs. NF controls which is not found in other forms of heart failure or HCM. It is not currently known why TTR accumulates in the myocardium during ATTR-CM. Our finding of increased endocytotic and vesicular trafficking proteins is surprising in that TTR deposition is toxic to the myocardium and limiting the uptake of TTR would likely be beneficial to cardiac function. This finding does provide a possible mechanism for ATTR-CM pathogenesis; however, this remains to be explored. Other upregulated proteins include cytoskeletal proteins, which have been observed in other HF syndromes and may provide insight into diastolic dysfunction in ATTR-CM independent of amyloid fibril deposition^12, 17, 18, 29^. Mitochondrial and protein homeostatic proteins were downregulated in ATTR-CM as were mitochondrial transcription/translation proteins. This too is seen in HFpEF, HFrEF, and HCM^12, 17, 18, 29^. Surprisingly, these proteins are downregulated to a greater extent than in any of the other syndromes. This may be a result of the tissue being extracted from explanted hearts from end-stage ATTR-CM patients. While true that the HFpEF tissue proteome used for comparison was obtained by endomyocardial biopsy, both the HCM and HFrEF proteomes were also taken from explanted hearts from end-stage patients^17^, so this cannot be the only explanation.

The finding that sarcomere proteins were downregulated is striking. The majority of heart failure syndromes and cardiomyopathies are associated with a pathologic hypertrophy, whether eccentric or concentric, and are often associated with upregulation of sarcomere proteins^12, 17, 18, 29^. Notably, proteomic studies from patients with cardiac cachexia also showed an upregulation of sarcomere proteins, which further highlights the significance of this finding^30^. Apart from downregulation of sarcomere proteins from a genetic etiology to our knowledge, this is the only acquired cardiac syndrome where such a phenotype is observed. This is accompanied by no change in the ratio of β and Ill myosin heavy chain, which is often elevated in heart failure^24^. One possibility is a change in sarcomere protein isoform, as observed in HFrEF, but this was not seen. In fact, the ratio of β and Ill myosin heavy chain, which is often elevated in heart failure, was not different in ATTR-CM patients vs. control^24^. Another possibility for this observation is the presence of excess TTR within the protein fraction. Given that all patients have ATTR-CM, then, for the same amount of total protein, the absolute mass of cardiomyocyte proteins would be expected to be reduced. If this were the case, all sarcomere proteins would be downregulated as observed. However, we found that calcium handling proteins, like cardiac phospholamban, also unique to cardiomyocytes, was not different in ATTR-CM vs. NF patients, so this is unlikely the reason Restoration of sarcomere protein content may be one potential novel therapeutic target for future consideration.

One prior study of permeabilized tissue muscle bundles from ATTR-CM patients showed a decrease in myocyte contractility as well as increased sensitivity to calcium and impaired relaxation^23^. The mechanism for this is not known, although the current study may provide insights. Reduced total myosin content limits the ability of the myocytes to participate in crossbridge formation, which may lead to impaired contractility and systolic dysfunction in ATTR-CM. Post-translational modifications of the sarcomere is another significant regulator of myocyte function. Our proteomics analyses identified ROCK-II, a kinase known to reduce myocyte contractility and increase calcium sensitivity in vivo, to be significantly upregulated in ATTR-CM which may also contribute to systolic impairment^26^. The accepted mechanism for ROCK-II mediated regulation of RLC phosphorylation involves an increase in the phosphorylation of myosin light chain phosphatase, which reduces its activity, ultimately leading to increased RLC phosphorylation^31-34^. The reduction in contractility is often attributed to increased S19 phosphorylation, but this was not observed^31-34^. However, increased RLC S15 phosphorylation was observed and is known to increase myocyte contractility by increasing calcium sensitivity and slowing down myosin detachment^35^. Increased RLC S15 phosphorylation may serve as a compensatory mechanism to ameliorate reduced contractility. However, while this may be beneficial under normal circumstances, this would be pathologic in ATTR-CM, which would serve to further increase calcium sensitivity and worsen diastolic function.

New therapeutic strategies have greatly improved the prognosis for ATTR-CM patients; however critical limitations remain. To date, management of cardiac function is supportive and is often restricted preload and afterload optimization. This is complicated by difficulties associated with early detection of ATTR-CM from lack of a genomic indicator, ultimately yielding delayed clinical manifestation of cardiac dysfunction and few options for intervention^34^.

TTR stabilizers (e.g. tafamidis) have been effective at slowing disease progression by stabilizing the tetrameric state of circulating TTR preventing misfolding and amyloid fibril formation, having shown the most benefit when ATTR-CM is detected early^36-39^. Furthermore, TTR silencers are increasing in prevalence as an option for addressing hereditary ATTR-CM, lifetime administration of TTR stabilizers coupled with standard heart failure intervention remains the sole standard therapeutic intervention currently available for chronic treatment of ATTRwt-CM^35^. Importantly, use of the medications does not reverse the course of disease but rather slows its progression^35–38^. There are currently no approved treatments capable of improving cardiac function or specifically targeting the dysfunctional processes within the myocardium. This in-depth proteomics analyses provides critical new insight into ATTRwt-CM by identifying the pathogenic pathways in the heart, allowing for the development of novel and targeted therapeutic interventions for ATTRwt-CM patients.

### Limitations

This study has several limitations. The sample size is low (n=4) for ATTR-CM patients, given the relatively low incidence of heart transplantation in ATTR-CM patients. This results in reduced power for detection of potential significant findings. TTR was not detected in our proteome, as the protein requires high resolution mass spectrometry for detection. The study was restricted ATTR-CM patients with wild type TTR, and relevance to ATTR-CM patients with variant TTR protein is unknown. All findings in the study are correlative and can only provide insight into potential mechanisms of disease. These are actively being explored.

## SOURCES OF FUNDING

Supported by NIH – National Heart Lung and Blood Institute grants HL169273 (MJR), Amyloidosis Research Foundation Donald C. Brockman Award (MJR), Pfizer Transthyretin Cardiac Amyloidosis Award #61667555, Maryland Stem Cell grant (2023-MSCRFL-6271), and an American Heart Association Career Development Award 18CDA34110140 and Transformational Project Award 20TPA35500008 (MJR). This material is based upon work supported by the National Science Foundation Graduate Research Fellowship under Grant No. (DGE2139757).

## ABBREVIATIONS

APO: apolipoprotein
ATTR-CM: amyloid transthyretin cardiomyopathy
HCM: hypertrophic cardiomyopathy
HFpEF: heart failure with a preserved ejection fraction
HFrEF: heart failure with a reduced ejection fraction
MS: mass spectrometry
TTR: transthyretin
WT: wild type

## Notes

### Competing Interest Statement

The authors have declared no competing interest.

